# Off-target expression of Cre-dependent adeno-associated viruses in wild type C57BL/6J mice

**DOI:** 10.1101/2021.09.08.459310

**Authors:** Justin J. Botterill, Abdessattar Khlaifia, Brandon J. Walters, Mark A. Brimble, Helen E. Scharfman, Maithe Arruda-Carvalho

**Affiliations:** Department of Psychology, University of Toronto Scarborough, Toronto, ON, M1C1A4, Canada; Department of Cell & Systems Biology, University of Toronto Mississauga, Mississauga, ON, L5L1C6, Canada; Department of Surgery, St. Jude Children’s Research Hospital, Memphis, TN, 38105, USA; Center for Dementia Research, The Nathan Kline Institute for Psychiatric Research, Orangeburg, NY, 10962, USA; Department of Child & Adolescent Psychiatry, Neuroscience & Physiology and Psychiatry and the New York University Neuroscience Institute, New York University Langone Health, New York, NY, 10016, USA; Department of Cell and Systems Biology, University of Toronto, Toronto, ON, M5S3G5, Canada

**Keywords:** Immunofluorescence, antibody amplification, double inverted open reading frame, fear conditioning, cFos, Cre/loxP, DREADDs

## Abstract

Adeno-associated viruses (AAVs) are a commonly used tool in neuroscience to efficiently label, trace, and/or manipulate neuronal populations. Highly specific targeting can be achieved through recombinase-dependent AAVs in combination with transgenic rodent lines that express Cre-recombinase in specific cell types. Visualization of viral expression is typically achieved through fluorescent reporter proteins (e.g., GFP or mCherry) packaged within the AAV genome. Although non-amplified fluorescence is usually sufficient to observe viral expression, immunohistochemical amplification of the fluorescent reporter is routinely used to improve viral visualization. In the present study, Cre-dependent AAVs were injected into the hippocampus and cortex of wild-type *C57BL/6J* mice. While we observed weak but consistent non-amplified off-target DIO expression in *C57BL/6J* mice, antibody amplification of the GFP or mCherry reporter revealed extensive Cre-independent viral expression. Off-target expression of DIO constructs in wild-type *C57BL/6J mice* occurred independent of vendor, AAV serotype or promoter. We also evaluated whether Cre-independent expression had functional effects via Designer Receptors Exclusively Activated by Designer Drugs (DREADDs). The DREADD agonist C21 had no effect on contextual fear conditioning or cFos expression in DIO-hM3Dq-mCherry+ cells of *C57BL/6J* mice. Taken together, our results indicate that DIO constructs have considerable off-target expression in wild type subjects. Our findings are particularly important for the design of experiments featuring sensitive systems and/or quantitative measurements that could be negatively impacted by off-target expression.

**Significance Statement:** Adeno-associated viruses (AAV) are widely used in neuroscience because of their safety and ease of use. Combined with specific promoters, Cre/loxP, and stereotaxic injections, highly specific targeting of cells and circuits within the brain can be achieved. In the present study we injected Cre-dependent AAVs into wild-type *C57BL/6J* mice and found considerable Cre-independent viral expression of AAVs encoding mCherry, GFP, or hM3Dq following immunohistochemical amplification of the fluorescent reporter protein. Importantly, we observed no functional effects of the Cre-independent expression in the hippocampus, as C21 had no detectable effect on DIO-hM3Dq-mCherry infected neurons in *C57BL/6J* mice. Given the widespread use of DIO rAAVs by the neuroscience community, our data supports careful consideration when using DIO constructs in control animals.

## Introduction

A main goal of neuroscience is to understand the roles of specific cell types and circuits underlying neurodevelopment, behavior, and disease. Adeno-associated virus (AAV) represents a powerful tool for neuroscientists to address these questions via labelling and manipulating cell types and circuits. AAV is a *dependoparvovirus* comprising a small 4.7kb single-stranded DNA genome with an unenveloped icosahedral capsid (Grieger and Samulski, 2005; Betley and Sternson, 2011; Haery et al., 2019; Haggerty et al., 2020). Recombinant AAVs (rAAVs) used in research and clinical applications are modified from wild-type (WT) AAVs and use an expression cassette to drive transgene expression. The rAAV expression cassette typically consists of a promoter, transgene, and polyadenylation signal flanked by inverted terminal repeats (ITRs) (Saunders and Sabatini, 2015). A major advantage of rAAVs is their durable transgene expression (months-years) and limited pathogenic profile (Naso et al., 2017; Haery et al., 2019; Haggerty et al., 2020).

The Cre/loxP system is a powerful site-specific recombinase used to insert, delete, or invert DNA sequences between loxP sites (Sauer and Henderson, 1988; Sengupta et al., 2017; Fischer et al., 2019). Using the Cre/loxP system, discrete cell populations can be targeted through a combination of transgenic mice and viral injections. Using this method, rodents are genetically modified to express Cre in specific cell types, and therefore the injection of Cre-dependent constructs should only recombine in Cre-expressing cells within the injected area. Double inverted open reading frame (DIO) constructs are a common method to achieve Cre-dependent activation of genes. DIO constructs rely on two pairs of recombination-incompatible lox sites (loxP and lox2722) that surround the transgene which is in the inverse orientation. However, in the presence of Cre, the DIO cassette is reverted, allowing expression of the transgene (Fenno et al., 2011). DIO cassettes are widely used because DIO is considered to have low off-target expression (Fischer et al., 2019) due to the transgene being in the incorrect orientation. Additionally, DIO is much smaller than other constructs with a similar goal, facilitating its use in AAVs.

Visualization of rAAV expression is typically achieved with fluorescent reporter proteins; either fused to a transgene of interest or inserted into its own reading frame (Smith et al., 2016). Fluorescent reporters exhibit relatively strong and permanent expression in transduced neurons and depending on the method employed can reveal expression in dendrites or axons (Betley and Sternson, 2011; Saleeba et al., 2019). The fluorescent reporter can also be inserted between loxP sites to allow for Cre-dependent expression of fluorescence signal (Betley and Sternson, 2011; Saunders and Sabatini, 2015; Saleeba et al., 2019). However, a limitation of fluorescent reporters is that expression can be weak in certain applications. For example, fluorescence can decline substantially following exposure to fixatives or high temperatures during tissue processing (Alkaabi et al., 2005). To circumvent weak rAAV fluorescence *ex vivo*, many studies amplify expression with antibodies against reporter proteins (e.g., GFP, mCherry) to improve visualization of fluorescence expression (Deverman et al., 2016; McGlinchey and Aston-Jones, 2018; Murata and Colonnese, 2020; Iwasaki and Ikegaya, 2021). Subjects that lack Cre are often used as controls for the behavioral or cellular effects of Cre-dependent viruses (Alexander et al., 2018; Bonaventura et al., 2019; Mahler et al., 2019), under the premise that these constructs limit expression to Cre-positive cells.

In the present study, we found consistent Cre-independent expression of DIO constructs in *C57BL/6J* mice injected across different brain regions. While Cre-dependent rAAVs showed minimal non-amplified fluorescence in brain sections of WT *C57BL/6J* mice, fluorescence signal amplification revealed numerous positive cells within the region of viral infection. To address whether the amplified fluorescence signal had functional effects, we utilized the Cre-dependent Designer Receptors Exclusively Activated by Designer Drugs (DREADDs) construct hM3Dq-mCherry, which is a modified human muscarinic M3 receptor that promotes neuronal excitation when activated (Roth, 2016). We found no detectable effect of the hM3Dq agonist C21 on fear behavior or immediate early gene activity in the hippocampus of WT *C57BL/6J* mice. Our results have important implications for the use of DIO constructs in control subjects, particularly in sensitive circuits or studies focusing on quantitative analyses such as cell counting or evaluating fluorescence signal.

## Materials and Methods

### Animals

Adult male and female mice aged 2-6 months were used for all experiments. For experiments testing Cre-dependent viral expression in mice lacking Cre-recombinase, we used WT *C57BL/6J* mice (Jackson Laboratory). Tyrosine hydroxylase-Cre (*TH-Cre, a kind gift from Dr. Jonathan Britt, McGill University*) (Lindeberg et al., 2004) and parvalbumin-Cre (*PV-Cre*, Jackson Laboratory) mice were used in a subset of experiments and genotyping for these lines was done in house using standard PCR protocols. Mice were bred in house and maintained on a 12hr light-dark cycle (lights on at 07:00h) with access to food and water *ad libitum*. Mice were housed in standard laboratory cages that contained corn cob bedding and a polycarbonate igloo shelter (Bio-Serv). Offspring were weaned with same-sex siblings on postnatal day 21 (2-5 mice per cage). All experiments were done during the light phase of the light-dark cycle. All animal procedures were approved by the Animal Care Committee at the [*Author University*]. Experimenters were blinded for all quantitative analyses.

### Stereotaxic surgery and viral injections

Mice underwent stereotaxic surgery between 2-5 months of age. Briefly, mice were injected intraperitoneally (i.p.) with a combination of ketamine (100 mg/kg) and xylazine (5 mg/kg) to induce anesthesia. Once anesthetized, the head was shaved and swabbed with iodine followed by 70% ethanol. Tear gel (Alcon) was applied to the eyes to prevent dehydration. Mice were then secured in a rodent stereotaxic apparatus (Stoelting) using ear bars. Body temperature was maintained throughout surgery with a heating blanket. An incision was made down the midline of the scalp using a scalpel, the connective tissue was excised, and then the skull was cleaned with sterile phosphate buffered saline (PBS, pH=7.4). An autoclaved cotton-tip applicator was briefly submerged in 30% hydrogen peroxide and gently applied to the skull surface to identify bregma. Using bregma as a reference point, craniotomies were made over the left medial prefrontal cortex (mPFC; +1.9mm anterior-posterior, 0.3mm medial-lateral), left anterior hippocampus (−2.1 mm anterior-posterior and -1.25 mm medial-lateral), left posterior hippocampus (−3.05mm anterior-posterior, -2.35mm medial-lateral), or ventral tegmental area (VTA; -3.15mm anterior-posterior, +/- 0.45mm medial-lateral). Experiments targeting the mPFC employed a single viral injection, whereas dual viral injections were administered for the hippocampus (anterior and posterior) and VTA (bilateral) experiments.

Virus was delivered using a 500nL Neuros Syringe (#65457-02, Hamilton Company) attached to the stereotaxic apparatus with a probe holder (#751873, Harvard Apparatus). The syringe was positioned above each craniotomy and the needle was lowered into the mPFC (−2.3mm below skull surface), hippocampus (−1.95mm anterior, -2.5mm posterior below skull surface) or ventral tegmental area (−4.5mm below skull surface). For each injection, 0.2µL of virus was injected at a rate of 0.06µL/minute. The following viral constructs were used: AAV5-EF1a-DIO-eYFP (≥4 × 10^12^vg/mL, UNC Core), AAV5-EF1a-DIO-mCherry (≥7 × 10^12^vg/mL, UNC Core), AAV8-hSyn-DIO-hM3D(Gq)-mCherry (≥5 × 10^12^vg/mL, UNC Core), or AAV5-hSyn-DIO-hM4D(Gi)-mCherry (≥8 × 10^12^vg/mL, Addgene #44362). The needle remained in place for an additional 5 minutes after each injection to allow for diffusion of the virus and then the needle was slowly removed from the brain. Ketoprofen (1 mg/kg, s.c.) was injected approximately 30 minutes prior to the end of surgery to reduce discomfort. The skull was cleaned with sterile PBS and the scalp was sutured with Vetbond tissue adhesive (3M). Mice were injected with 0.7mL of warmed physiological saline at the end of surgery to support hydration. Mice were then transferred into a clean cage located on a heating blanket. Mice were returned to their colony room once fully ambulatory. Ketoprofen (1 mg/kg, s.c.) was administered 24 and 48 hours after surgery to reduce post-surgical discomfort.

### Contextual fear conditioning

Mice received a post-surgical recovery of 2 weeks prior to behavioral testing. Mice were transferred to a dedicated procedures room and injected with Compound 21 (C21; 1.5mg/kg, 0.2mg/mL dissolved in 0.9% NaCl; HelloBio) one hour before fear training. Mice underwent contextual fear conditioning as previously described (Arruda-Carvalho et al., 2011; Guskjolen et al., 2018). Briefly, mice were individually placed in stainless steel fear conditioning apparatus (32cm wide, 25.5cm high, 25.5cm deep) that contained shock grid floors (36 rods, 2mm diameter). The fear conditioning apparatus was located inside a sound-attenuated chamber (63.5cm wide, 36.8cm high, 74.9cm deep, #NIR-022MD, Med Associates). A 2-minute acclimation period was used to assess baseline behavior. Foot-shocks (0.5mA, 2s duration) were delivered 120s, 180s, 240s, 300s, and 360s after mice were placed in the chamber. Mice remained in the chamber for 60s after the final foot-shock and were then returned to their home cage. Mice were returned to the colony housing room and left undisturbed until the context test on the following day. Contextual fear memory was assessed 24 hours after the training session. Mice were returned to the same fear conditioning chamber as the previous day in absence of foot-shocks and freezing behavior was evaluated over 8 minutes. Notably, C21 was not administered prior to testing.

Conditioned freezing was identified by the absence of movement except those necessary for respiration (Blanchard and Blanchard, 1972; Fanselow, 1980). Freezing behavior was scored automatically using the Med Associates VideoFreeze software.

### Perfusions and sectioning

Mice were euthanized 2-3 weeks after surgery to evaluate viral expression. Subjects were injected with Avertin (250mg/kg, i.p.) and once under deep anesthesia, transcardially perfused with 15 mL of room temperature saline, followed by 15 mL of cold 4 % paraformaldehyde (PFA). The brains were extracted and stored overnight at 4 °C in 4 % PFA. The brains were sectioned at 50 µm in the coronal plane (VT 1000, Leica) and stored at -20 °C in a cryoprotectant solution comprised of 60% glycerol and 0.01% sodium azide in 0.1M phosphate buffered saline (PBS).

### Immunofluorescence

Immunofluorescence staining was performed on free floating sections. Sections were washed in 0.1M PBS (3 × 5 min each) and then incubated in blocking solution comprised of 5% normal goat serum and 0.25% Triton X-100 in 0.1M PBS for 30 min. Amplification of the viral signal was achieved by incubating sections with polyclonal rabbit anti-mCherry (1:2000, #ab167453, Abcam, RRID: AB_2571870) or polyclonal chicken anti-GFP (1:2000, #ab13970; Abcam, RRID: AB_300798) primary antibodies diluted in blocking solution. Sections were incubated with the primary antibodies overnight at 4 °C on a rotary shaker under gentle agitation. On the following morning, sections were incubated in goat anti-rabbit Alexa 568 (1:500, #A11011, ThermoFisher, RRID: AB_143157) or goat anti-chicken Alexa 488 (1:500, #A11039, ThermoFisher, RRID: AB_2534096) secondary antibodies for 2 hours. Sections were then counterstained with Hoechst 33342 (1:2000 diluted in 0.1M PBS; ThermoFisher). Sections were then rinsed in 0.1M PBS, mounted onto gelatin-coated slides, air dried for 30 min, and coverslipped with Citifluor anti-fade mounting medium (#17970, Electron Microscopy Sciences).

A subset of tissue was processed for mCherry and cFos using tyramide signal amplification (TSA). Briefly, sections were rinsed in 0.1M PBS, followed by 1% H_2_0_2_ in 0.1M PBS to quench endogenous peroxidase activity. Sections were then incubated overnight at 4 °C with polyclonal rabbit anti-cFos (#226 003, Synaptic Systems, RRID: AB_2231974) and monoclonal rat anti-mCherry (#M11217, ThermoFisher, RRID: AB_2536611) primary antibodies in 0.1M Tris-buffered saline containing 0.5% Roche Blocking Reagent (#11096176001; Sigma). On the following day, sections were incubated in goat anti-rat Alexa 568 secondary antibody (1:500, #A11077, ThermoFisher, RRID: AB_2534121) and donkey anti-rabbit horseradish peroxidase conjugated secondary antibody (1:500, #711-036-152; Jackson ImmunoResearch Laboratories, RRID: AB_2340590) for 1 hour each. Next, TSA was performed using fluorescein tyramide (1:100) diluted in 0.1M borate buffer containing 0.01% H_2_0_2_ solution. Sections were counterstained with Hoechst 33342 (1:2000), mounted onto slides, air dried, and coverslipped with anti-fade mounting medium as described above.

### Diaminobenzidine tetrahydrochloride (DAB) staining

Immunohistochemistry for brightfield microscopy was performed using standard protocols (Koshimizu et al., 2021). Sections were rinsed in 0.1M PBS, endogenous peroxidase activity was quenched with 0.3% H_2_0_2_, and blocked in 5% normal goat serum. Sections were incubated with polyclonal rabbit anti-mCherry primary antibody (1:8000) diluted in blocking solution overnight at 4 °C. On the following day sections were incubated in biotinylated goat anti-rabbit secondary antibody (1:500, #BA-1000, Vector laboratories, RRID: AB_2313606) and avidin-biotin complex (1:500, PK-6100, Vector laboratories, RRID: AB_2336819). Immunoreactivity was visualized by incubating sections in 0.5mg/mL 3,3’-diaminobenzidine tetrahydrochloride (Sigma), 40µg/mL ammonium chloride (Sigma), 25mg/mL (D+)-glucose (Sigma), and 3µg/mL glucose oxidase (Sigma) for approximately 5 minutes. Sections were mounted onto gelatin-coated slides and allowed to dry overnight. Sections were then dehydrated using a graded alcohol series (70, 95, 100%), cleared with xylenes, and coverslipped with Permount mounting medium (Electron Microscopy Sciences).

### Image acquisition

Images were acquired with a Nikon Eclipse Ni-U epifluorescence microscope running NIS-elements software (v. 5.11.03, Nikon). Immunofluorescence was visualized with an LED illumination system (X-Cite 120 LED Boost, Excelitas Technologies) and captured with a Nikon DS-Qi2 digital camera. Immunofluorescence images were acquired using Plan Fluor 4x, Plan-Apochromat 10x DIC N1 or Plan Fluor 20x DIC N2 objectives. Brightfield images were acquired with a 10x objective on an Olympus BX61 microscope. Figures were made using Adobe Photoshop 22.5. When brightness and/or contrast adjustments were made in a figure, these changes were made equally to all photomicrographs.

### Quantification

Cell counts were done manually using ImageJ software (v. 1.53e) by experimenters blinded to treatment conditions. For cell counts in the hippocampus, counts were performed on a minimum of 5 sections per subject that spanned the rostral-caudal extent of the hippocampus. For cell counts in the mPFC, approximately 3-4 sections were counted per subject due to the smaller number of available sections for this region. Cell counts were performed for both the injected and non-injected hemispheres for each subject. The average number of cells per section was calculated by summing the total number of cells counted in the injected or non-injected hemisphere and dividing by the number of sections that were analyzed.

### Statistical analysis

All results are presented as the mean ± standard error of the mean (SEM). Statistical comparisons were made using Prism 9.0 (GraphPad) with statistical significance (*p*< 0.05) denoted on all graphs with an asterisk. Comparisons of independent groups were made using two-tailed unpaired t-tests. Two-way repeated-measures ANOVAs were used to analyze parametric data with multiple comparisons followed by Tukey’s post hoc test with corrections for multiple comparisons when appropriate. Normality of parametric data sets were confirmed by the D’Agostino & Pearson normality test (Prism 9.0). Non-parametric data sets were analyzed with a Mann-Whitney U test. Potential sex differences were examined for each data set and indicated no significant differences between male and female mice (all p values >0.154). Male and female mice were therefore pooled for each dataset, but for transparency, all graphs show individual data points for male (dotted) and female (clear) mice.

## Results

### Fluorescence signal amplification of DIO constructs in Cre-positive and WT *C57BL/6J* mice

First, we evaluated non-amplified and amplified fluorescence of a DIO-mCherry construct in the *TH-Cre* mouse line, which labels dopaminergic neurons in midbrain structures such as the VTA (Lammel et al., 2015; Popescu et al., 2016). AAV5-EF1a-DIO-mCherry was injected bilaterally into the VTA of *TH-Cre* mice (**Figure 1A**). Near the injection site, non-amplified and amplified fluorescence showed a pattern of fluorescence consistent with previous reports (Lammel et al., 2015), but mCherry amplification produced a substantial increase in fluorescence signal. We also evaluated long-range projections from the VTA to nucleus accumbens (NAc), dorsal striatum (DS), and mPFC. The NAc and DS showed a moderate amount of non-amplified fluorescence, whereas mCherry+ terminals in the mPFC were only slightly greater than background fluorescence. In contrast, amplifying the fluorescent signal revealed bright fluorescence in the NAc-DS and numerous mCherry+ terminals in the mPFC (**Figure 1B-1C**), in a pattern consistent with previous reports (Stuber et al., 2010; Lammel et al., 2015; Popescu et al., 2016; Ellwood et al., 2017).

**Figure 1.**
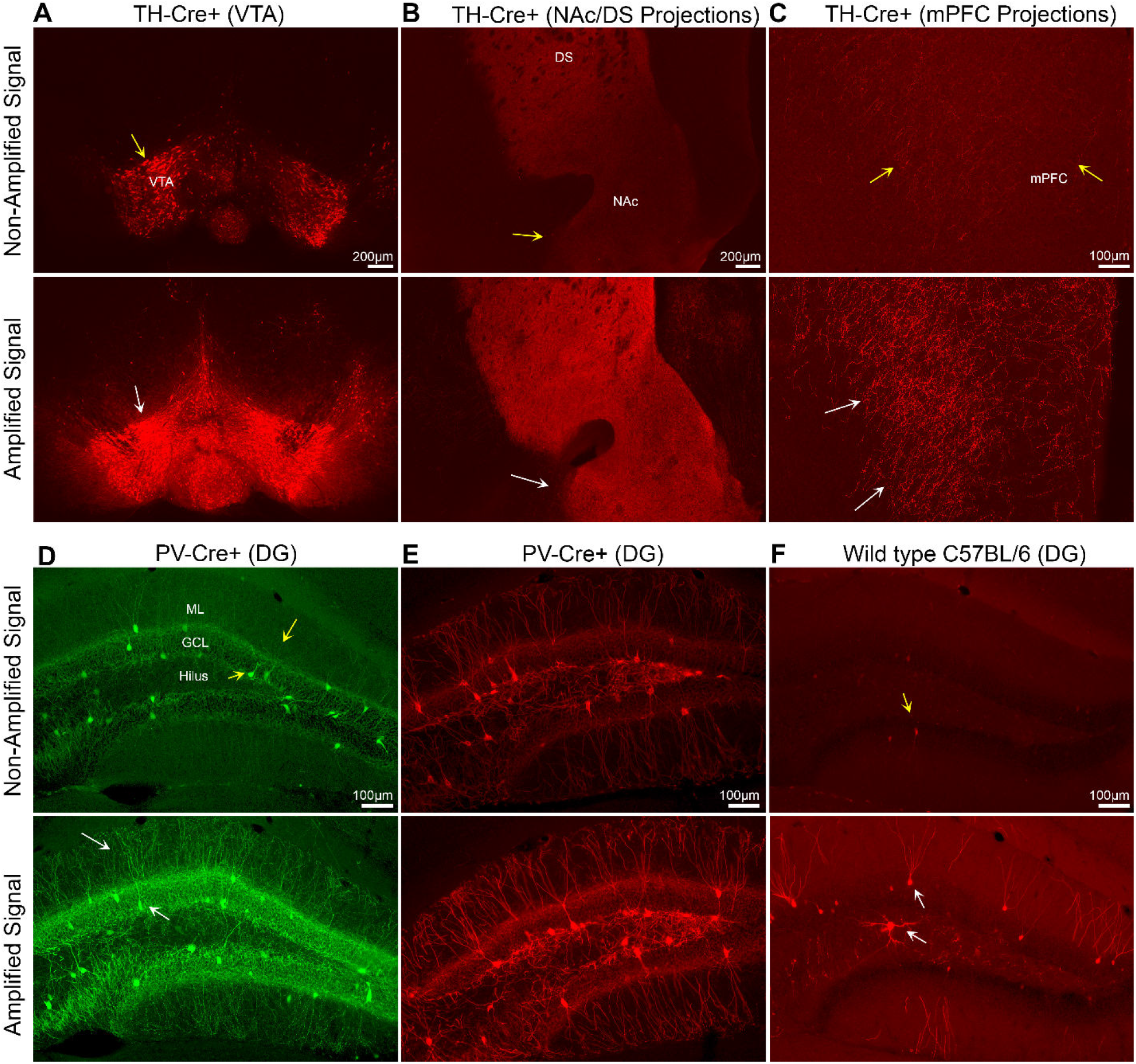
Antibody amplification of Cre-dependent viral expression. **(A)** Representative images from a *TH-Cre* mouse injected in the VTA with AAV5-EF1a-DIO-mCherry show a similar pattern of expression between non-amplified and amplified fluorescence (yellow and white arrows) **(B)** Long-range VTA to NAc/DS projections are considerably easier to visualize following mCherry amplification (yellow vs white arrow). **(C)** Similarly, non-amplified fluorescence of VTA to mPFC projections was generally weak (yellow arrows) and the fluorescence signal was significantly improved following mCherry amplification (white arrows). **(D-E)** Representative images from *PV-Cre* mice injected with **(D)** AAV5-EF1a-DIO-EYFP or **(E)** AAV5-EF1a-DIO-mCherry. The non-amplified fluorescence signal was similar between eYFP and mCherry constructs. Moreover, fluorescence signal amplification is similar to the non-amplified signal (yellow arrows) but is considerably brighter and it is easier to visualize (white arrows), especially the dendrites in the ML. **(F)** Representative images from a *C57BL/6J* mouse injected with AAV5-EF1a-DIO-mCherry show minimal non-amplified fluorescence (yellow arrow). Remarkably, amplification of adjacent sections from the same mouse revealed considerable mCherry expression within the DG (white arrows). VTA: ventral tegmental area, NAc: nucleus accumbens, DS: dorsal striatum, mPFC: medial prefrontal cortex, ML: molecular layer, GCL: granule cell layer. Scale bars: 200µm (4x objective), 100µm (10x objective).

We further compared non-amplified and amplified fluorescence in the hippocampus, a brain region widely studied and often targeted with DIO constructs. Using *PV-Cre* mice, which express Cre in parvalbumin interneurons, we targeted the dentate gyrus (DG) subfield of the hippocampus due to its well documented PV expression (Freund and Buzsaki, 1996; Pelkey et al., 2017). *PV-Cre* mice were injected with AAV5-EF1a-DIO-eYFP or AAV5-EF1a-mCherry (**Figure 1D-E**). In both cases, the non-amplified signal in the DG was characterized by bright fluorescence in somata and weaker fluorescence in fine processes, consistent with the overall patterns of parvalbumin immunoreactivity reported previously (Zou et al., 2016; Foggetti et al., 2019). Antibody amplification of GFP or mCherry resulted in brighter immunofluorescence signal, especially in fine processes, such as dendrites extending into the molecular layer (ML; **Figure 1D-E**). The results of the *TH-Cre*-positive and *PV-Cre*-positive experiments suggest that fluorescence signal amplification produces immunofluorescence expression that is faithful to non-amplified viral expression, but advantageous for visualizing cells or terminals with weak fluorescence.

Control experiments were also performed where AAV5-EF1a-DIO-mCherry was injected into the DG of WT *C57BL/6J* mice. Compared to the substantial fluorescence signal observed in the DG of Cre-positive mice, we observed minimal non-amplified fluorescence in control mice (**Figure 1F**). This observation is consistent with the requirement of Cre-recombinase for transgene expression and low “leak” with DIO constructs (Schnutgen et al., 2003; Atasoy et al., 2008; Saunders and Sabatini, 2015). However, amplification of DIO-mCherry revealed substantial immunofluorescence within the DG of control *C57BL/6J* mice (**Figure 1F**). The majority of amplified mCherry+ cells appeared to be granule cells (GCs), which reside in the principal cell layer of the DG known as the granule cell layer (GCL) and extend dendrites into the ML. We also observed sparse labeling of mCherry+ boutons in the hilus, consistent with expression of mCherry in dentate GC mossy fibers. Sparse labeling of large hilar cells was also observed. These data show that fluorescence signal amplification revealed substantial off-target expression in mice lacking Cre-recombinase.

### Non-amplified expression of DIO constructs in WT C57BL/6J mice

Next, we evaluated non-amplified fluorescence of DIO constructs in *C57BL/6J* mice to gain a better understanding of the off-target expression observed following fluorescence amplification. Non-amplified sections of *C57BL/6J* mice injected with AAV5-EF1a-DIO-eYFP or AAV5-EF1a-DIO-mCherry showed very few bright GFP+ or mCherry+ cells, respectively (**Figure 2**). This finding is consistent with the notion that Cre is required to drive transgene expression, but Cre-independent expression is possible (Fischer et al., 2019). Specifically, commercial vendors warn that recombination of loxP sites may occur during DNA amplification and viral production and result in Cre-independent transgene expression. However, this is thought to occur in a small number of viral particles (e.g., <1%) and therefore represent a minor source of off-target expression. Indeed, the few cells with bright fluorescence cannot explain the numerous cells we observed following fluorescence amplification. We found that increasing the exposure time and using higher power objectives (e.g., 20x) revealed numerous cells with weak fluorescence primarily restricted to the soma (**Figure 2, see insets**). Importantly, cells with weak fluorescence were only observed in the injected hemisphere. We hypothesize that these numerous but weakly labeled cells express low levels of the viral transgene (e.g., GFP or mCherry) and become strongly labeled following fluorescence signal amplification.

**Figure 2.**
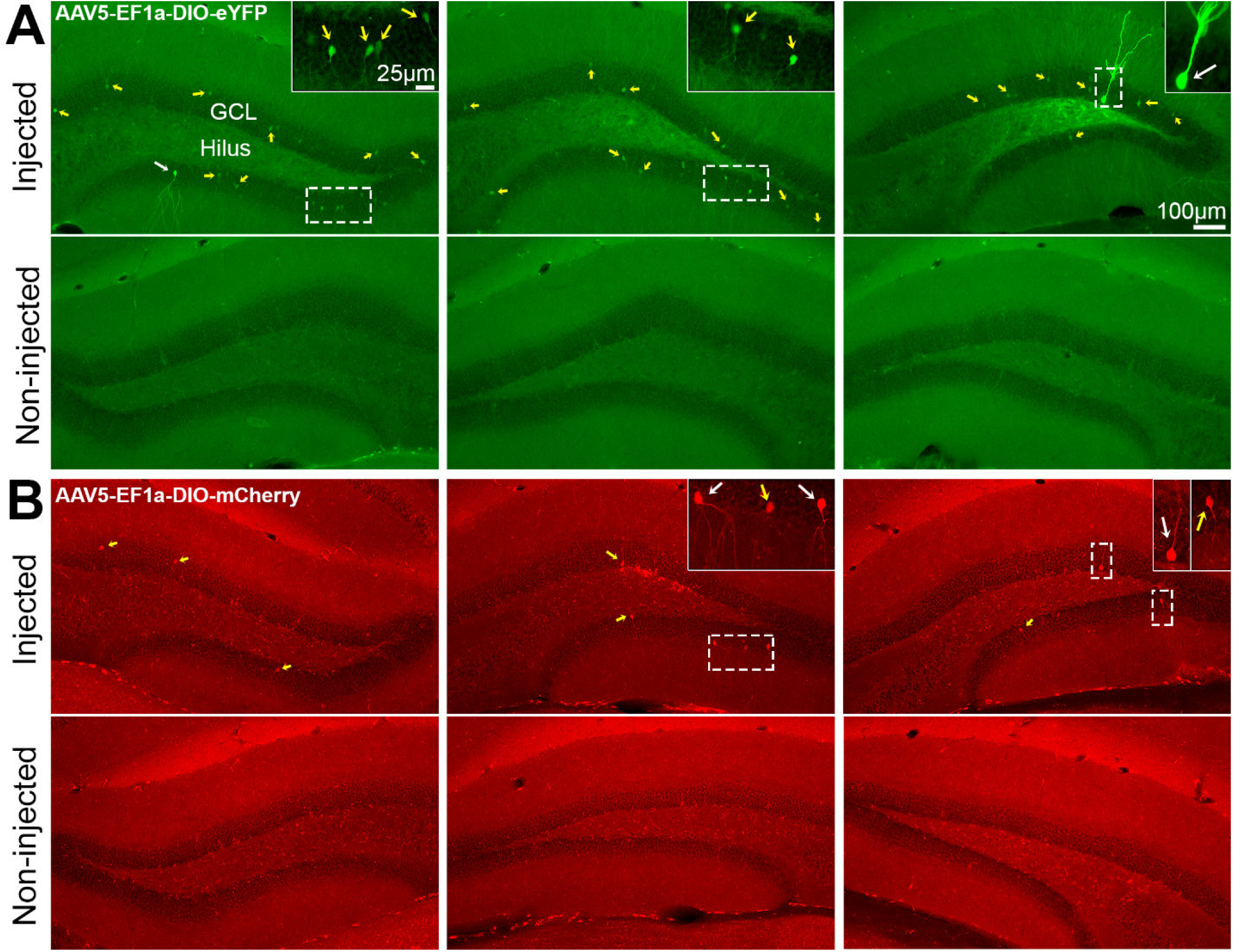
Non-amplified fluorescence of DIO constructs in WT C57BL/6J mice. Representative photomicrographs of non-amplified fluorescence signal in *C57BL/6J* mice injected with **(A)** AAV5-EF1a-DIO-eYFP or **(B)** AAV5-EF1a-DIO-mCherry. Non-amplified immunofluorescence was generally weak and primarily restricted to the soma (yellow arrows, see insets) of the injected hemisphere only. We hypothesize that the weak non-amplified immunofluorescence in these cells is significantly enhanced after antibody amplification. In addition, a very small number of cells with bright immunofluorescence throughout the cell body and its processes were observed (white arrows, see insets). Scale bars: 100µm (10x objective), Insets 25µm (20x objective).

### Fluorescence signal amplification of AAV5-EF1a-DIO-mCherry in WT *C57BL/6J* mice

To further investigate the off-target expression of AAV5-EF1a-DIO-mCherry in *C57BL/6J* mice, we quantified the number of mCherry+ cells in the anterior and posterior DG following fluorescence signal amplification (*n*=8; **Figure 3A-B**). Remarkably, amplified mCherry+ cells were found throughout the DG of *C57BL/6J* mice injected with DIO-mCherry (**Figure 3C**), almost exclusively restricted to the injected hemisphere (11.43 ± 1.40 cells/section, compared to the non-injected hemisphere 0.05 ± 0.02 cells/section; *Mann-Whitney U*=0, *p*<0.001; **Figure 3D**). Importantly, off-target expression was observed in all mice (*n*=8; range: 7.25 to 17.38 cells/section). We also processed a subset of sections with DAB and found that mCherry immunoreactivity was similar to the pattern of amplified DIO-mCherry immunofluorescence (**Figure 3-1**), indicating that our results were not attributable to non-specific fluorescence signal. These findings indicate that the off-target expression of DIO constructs in *C57BL/6J* mice revealed by amplification was highly reproducible.

**Figure 3.**
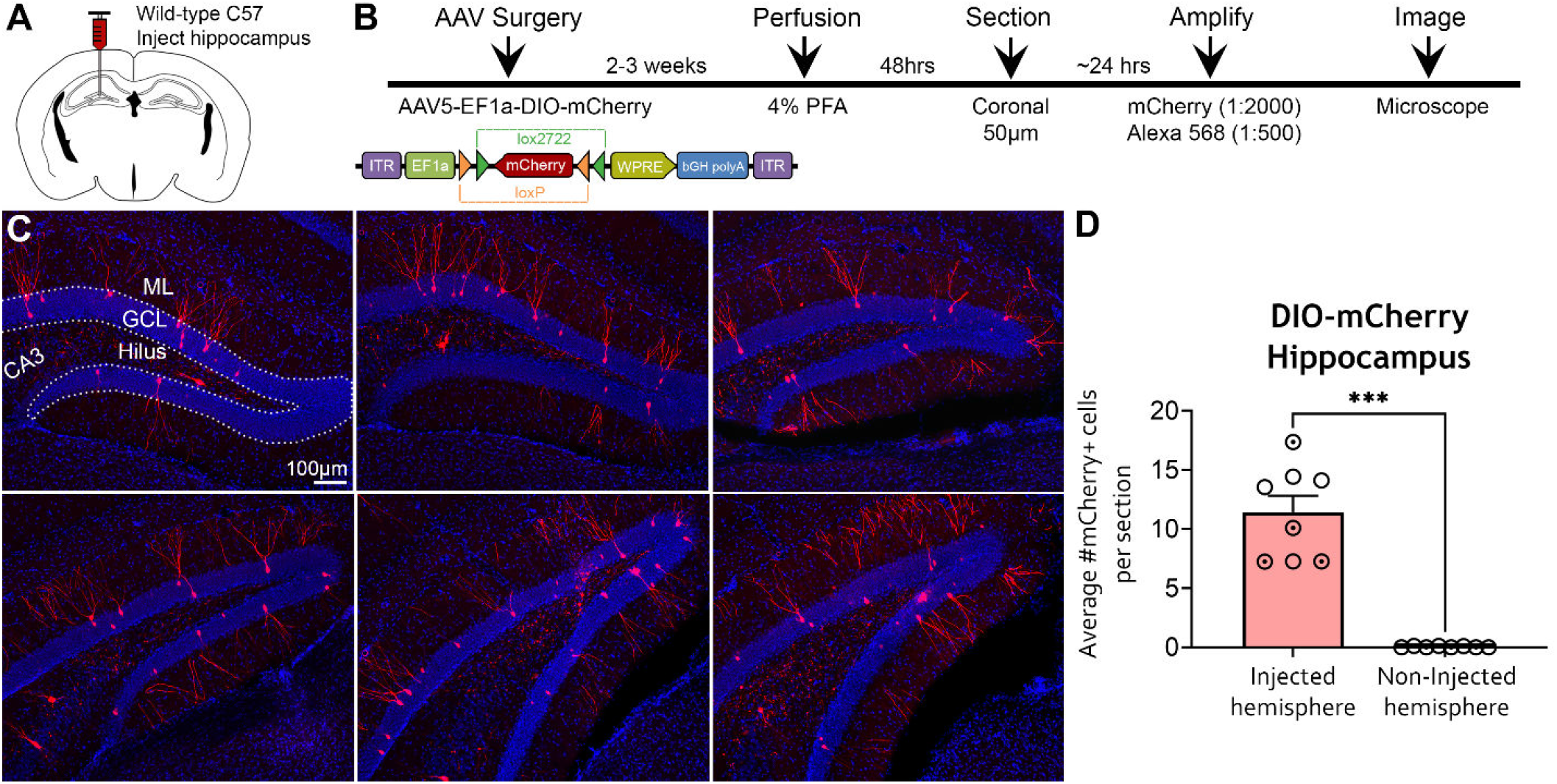
Amplified expression of DIO-mCherry in the hippocampus of WT *C57BL/6J* mice. **(A-B)** Experimental design and timeline. AAV5-EF1a-DIO-mCherry was injected into the anterior and posterior hippocampus of *C57BL/6J* mice (*n*=8) and perfused 2-3 weeks later. Brains were sectioned in the coronal plane and viral signal was amplified with rabbit anti-mCherry and goat anti-rabbit 568 antibodies. **(C)** Representative immunofluorescence of mCherry throughout the relatively dorsal (top panel) and caudal (bottom panel) DG. Expression of mCherry was primarily observed in the GCL and dendrites extending into the ML (putative dentate GCs). The amplified mCherry signal also resulted in labeling of mossy fibers and cells in the hilus. **(D)** Quantification of mCherry+ cells indicated that somatic expression was restricted to the injected hemisphere. Female (clear circles) and male (dotted circles) data points are identified, but no sex differences were found. GCL: granule cell layer, ML: molecular layer. ***p<0.001. Scale bar: 100µm.

**Figure 3-1.**
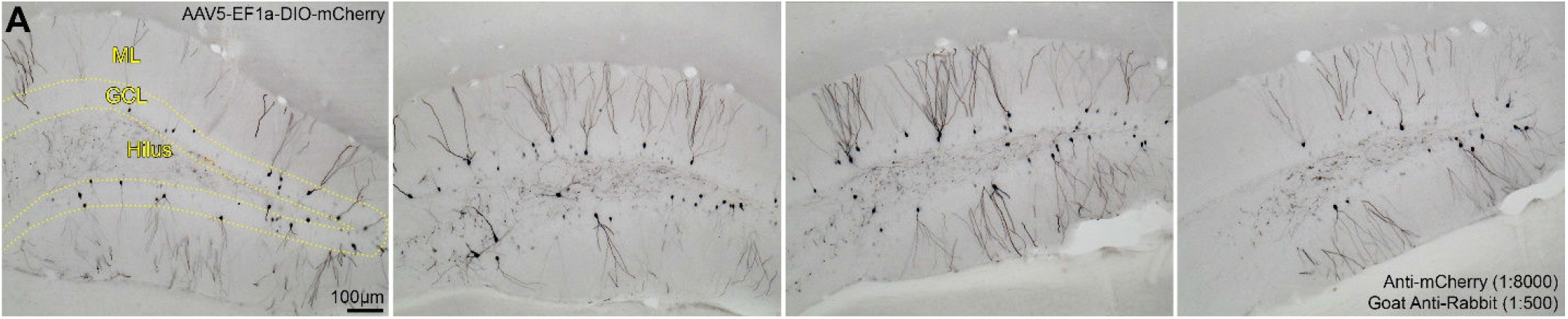
mCherry immunoreactivity in WT *C57BL/6J* mice injected with AAV5-EF1a-DIO-mCherry (A) Representative photomicrographs of mCherry immunoreactivity in *C57BL/6J* mice injected with AAV5-EF1a-DIO-mCherry. Overall, mCherry immunoreactivity was comparable to the pattern of expression observed with amplified DIO-mCherry immunofluorescence (see **Figure 3**). GCL: granule cell layer, ML: molecular layer. Scale bar: 100µm.

### Fluorescence signal amplification of AAV5-EF1a-DIO-eYFP in WT *C57BL/6J* mice

The high expression of amplified DIO-mCherry in *C57BL/6J* mice prompted us to investigate amplified expression using other DIO constructs. *C57BL/6J* mice (*n*=6) received injections of AAV5-EF1a-DIO-eYFP in the left anterior and posterior DG using identical parameters as the DIO-mCherry experiments (**Figure 4A**). Mice were euthanized 2-3 weeks after surgery and brains were sectioned and amplified with anti-GFP antibodies (**Figure 4B**). Amplification of GFP produced immunofluorescence in the DG that was more extensive than the DIO-mCherry experiments but shared a similar pattern (**Figure 4C**). Specifically, relatively sparse labeling of GFP+ cells was observed in the GCL similar to mCherry amplification. However, amplified GFP+ immunofluorescence resulted in robust labeling of dendrites in ML, compared to the relatively sparse labeling of the ML following mCherry amplification (**Figure 4C**). Furthermore, GFP+ immunofluorescence was more pronounced in the hilus, with expression observed in hilar cells and mossy fibers (**Figure 4C**). As with mCherry, GFP cell counts throughout the DG found that GFP+ cells were exclusive to the injected hemisphere (19.54 ± 3.46 cells, non-injected hemisphere: 0.00 ± 0.00 cells; *Mann-Whitney U*=0, *p*<0.001; **Figure 4D**). Taken together, these results demonstrate a highly specific pattern of amplified fluorescence signal of DIO constructs in *C57BL/6J* mice, independent of the construct used (DIO-mCherry or DIO-eYFP, **Figure 4-1**).

**Figure 4.**
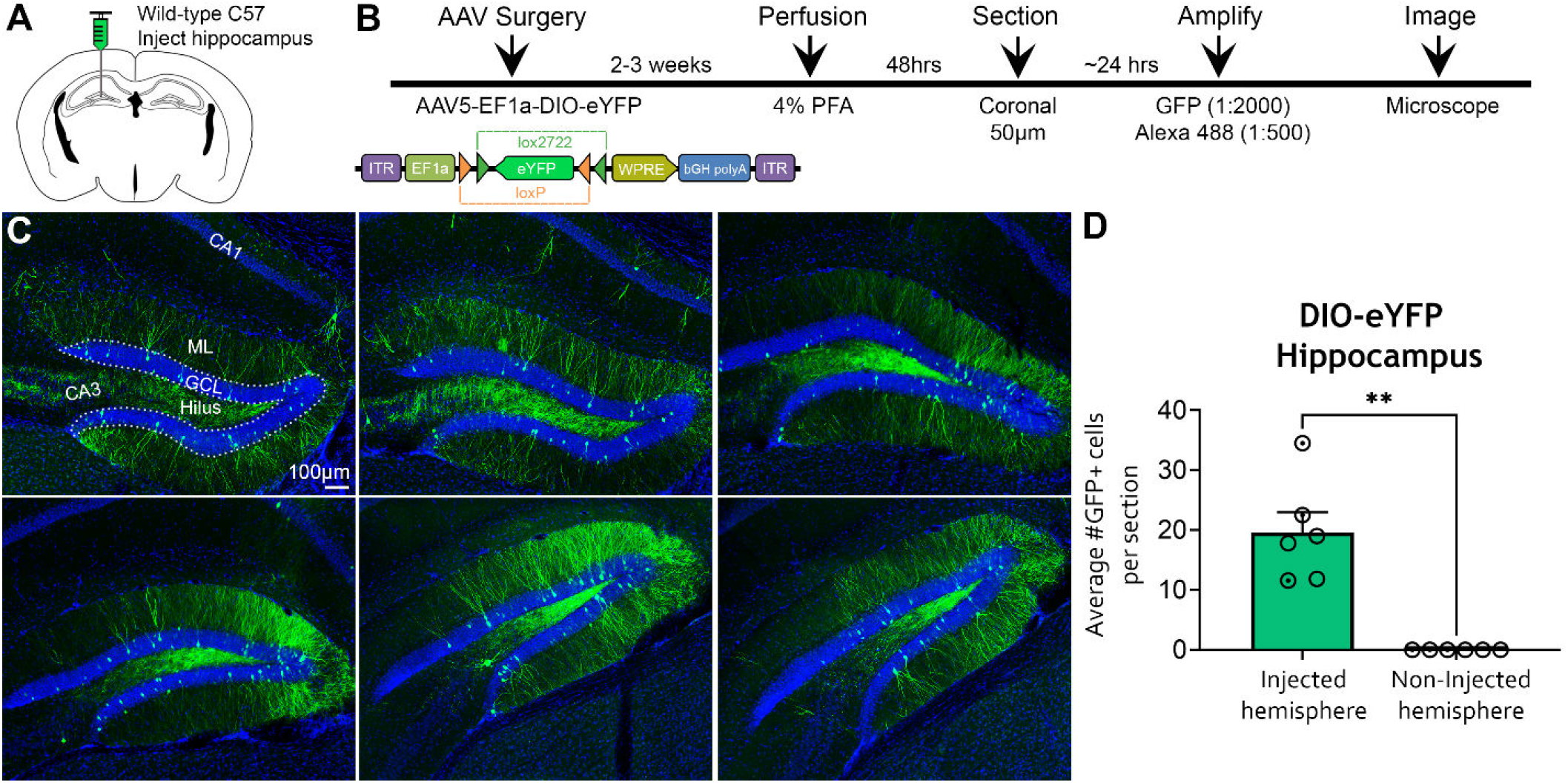
Amplified expression of DIO-eYFP in the hippocampus of WT *C57BL/6J* mice. **(A-B)** Experimental design and timeline. AAV5-EF1a-DIO-eYFP was injected into the anterior and posterior hippocampus of *C57BL/6J* mice (*n*=6) and perfused 2-3 weeks later. The eYFP signal was amplified with chicken anti-GFP and goat anti-chicken 488 antibodies. **(C)** Representative immunofluorescence of GFP throughout the DG. GFP expression was observed primarily in the DG, characterized by robust labeling of putative GCs within the GCL and their dendrites. The hilus also showed bright GFP signal, with expression in mossy fibers and hilar cells. **(D)** Quantification of GFP+ cells revealed that somatic expression was restricted to the injected hemisphere. Female (clear circles) and male (dotted circles) data points are identified, but no sex differences were found. GCL: granule cell layer, ML: molecular layer. **p<0.005. Scale bar: 100µm.

**Figure 4-1.**
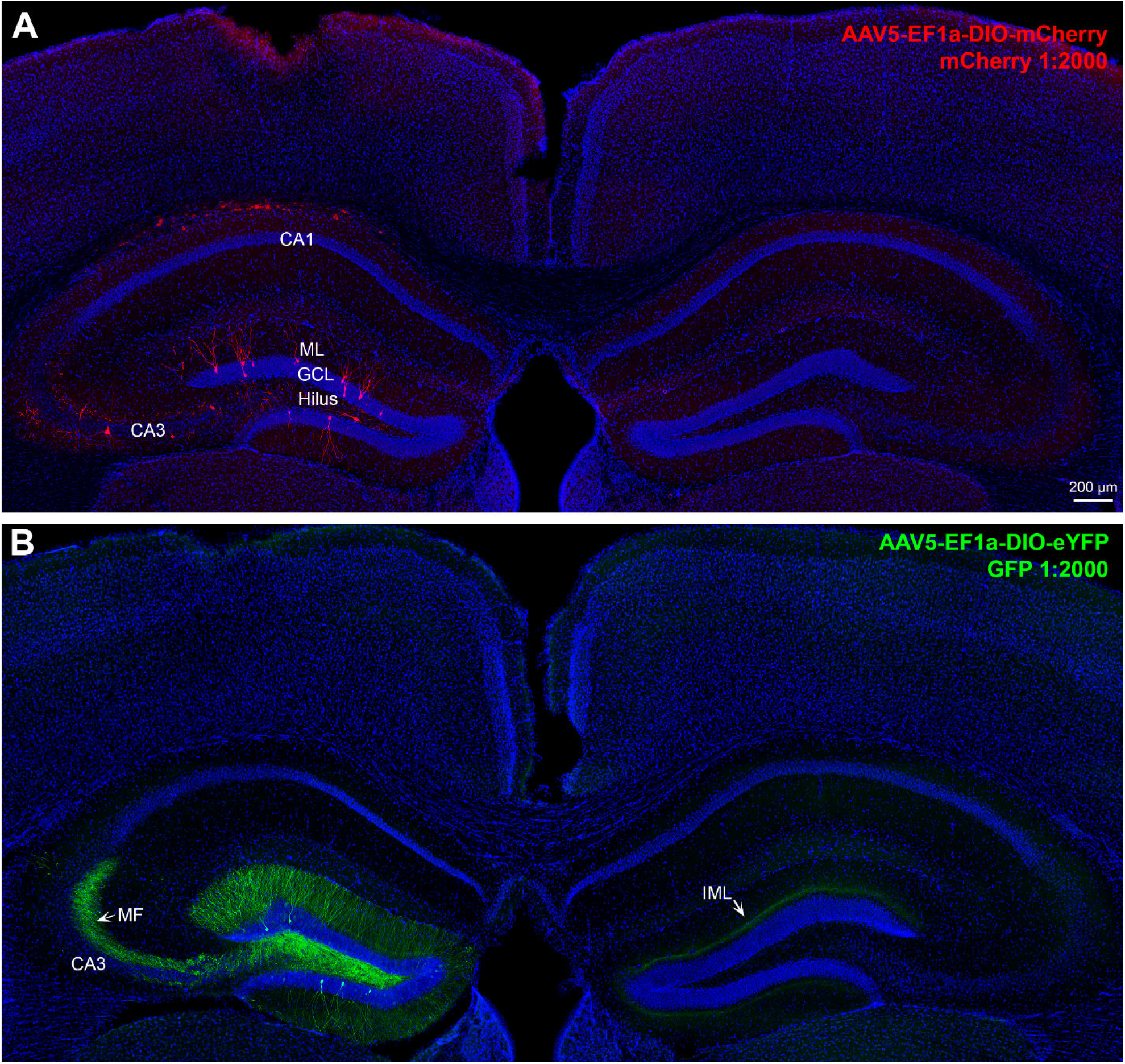
Fluorescence signal amplification of DIO-mCherry and DIO-eYFP is highly specific to the injection site in WT *C57BL/6J* mice. **(A)** Tile-scan of a *C57BL/6J* mouse injected with AAV5-EF1a-DIO-mCherry. Viral expression was amplified with mCherry antibody. The indent on the top of the left cortex represents drilling artifact near the injection site. The mCherry expression is primarily restricted to the injected (left) hippocampus, with mCherry+ cells observed in the GCL of the DG. There is also sparse labeling of mCherry+ cells in the CA3. **(B)** Tile-scan of a *C57BL/6J* mouse injected with AAV5-EF1a-DIO-eYFP. Viral expression was amplified with GFP and observed primarily within the injected (left) DG. Furthermore, GFP+ mossy fiber (MF) axons from dentate GCs were observed projecting to area CA3. Interestingly, commissural GFP+ axons, presumably from mossy cells, were observed within the IML of the contralateral hemisphere. Notably, there were no mCherry+ or GFP+ cells in the non-injected hemisphere. This result indicates that amplified fluorescence signal is highly specific to the target region and the projections of labeled cells. Scale bar: 200µm.

The amplified expression of mCherry and eYFP in the DG of *C57BL/6J* mice injected with Cre-dependent constructs led us to question whether off-target expression was unique to the DG or a general consequence of viral injections regardless of the region that was targeted. Serendipitously, we observed amplified immunofluorescence in hippocampal areas CA1 and/or CA2 when viral injections did not target the DG correctly (**Figure 4-2**).

**Figure 4-2.**
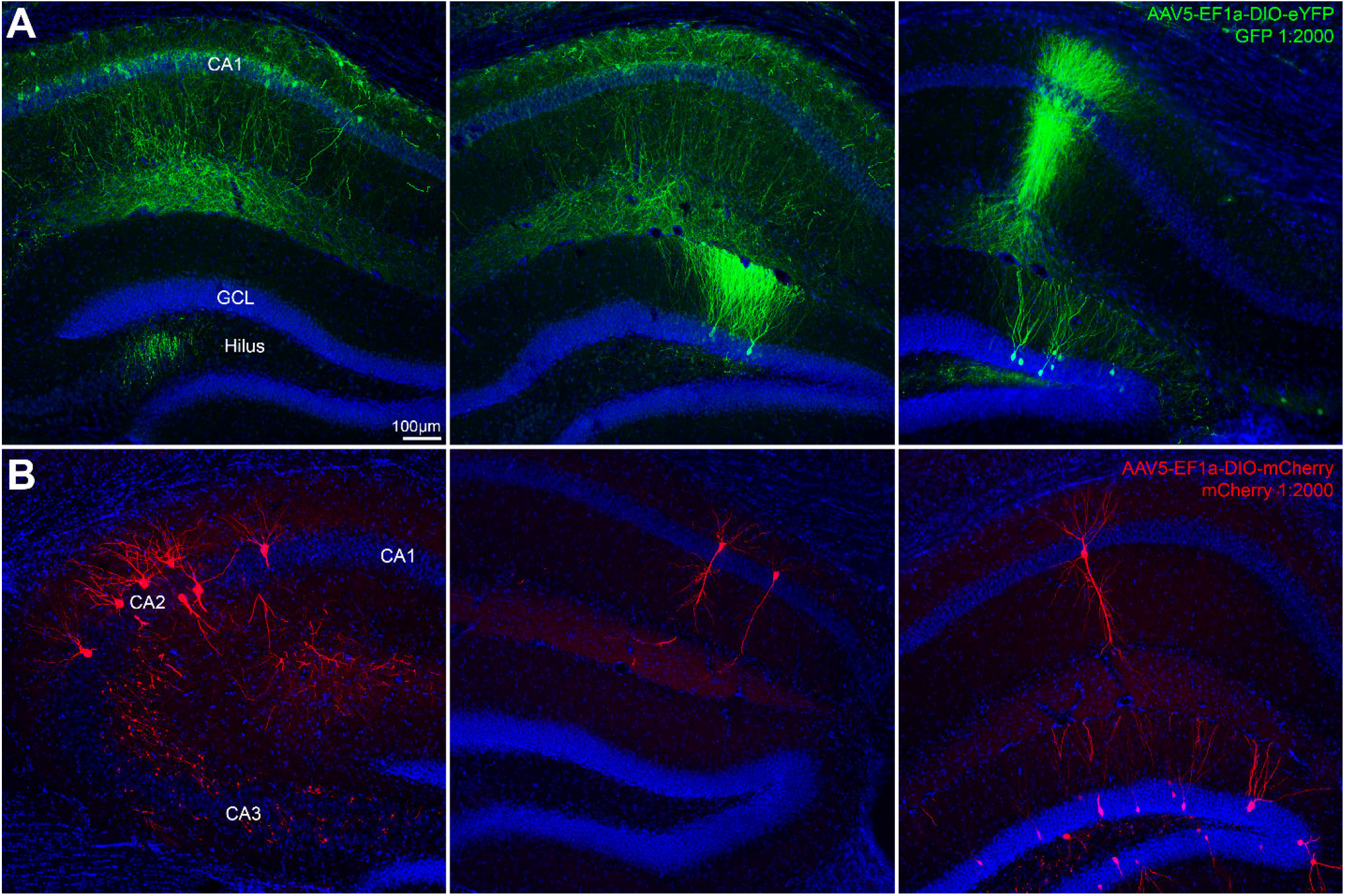
Fluorescence signal amplification in other subfields of the hippocampus. **(A-B)** Viral injections aimed at the DG occasionally resulted in mistargeting which led to amplified fluorescence signal in other subfields of the hippocampus, such as CA1 or CA2. This finding suggests that amplified viral expression was not unique to the DG, but rather specific to the injection site. GCL: granule cell layer. Scale bar: 100µm.

In addition, we specifically targeted the mPFC in *C57BL/6J* mice (*n*=6) using Cre-dependent eYFP (AAV5-EF1a-DIO-eYFP; **Figure 5A**). The experimental timeline for mPFC experiments was identical to the that of eYFP hippocampal injections (**Figure 5B**). Amplified GFP immunofluorescence was also observed in the mPFC (**Figure 5C**), at a similar rate as seen in DG (14.26 ± 3.29 cells/section in the injected hemisphere compared to 1.84 ± 1.20 cells/section in the non-injected hemisphere; *Mann-Whitney U*=1, *p*=0.004; **Figure 5D**). Overall, these findings suggest that Cre-independent, DIO construct expression is specific to the viral injection site, and not tied to a particular brain region.

**Figure 5.**
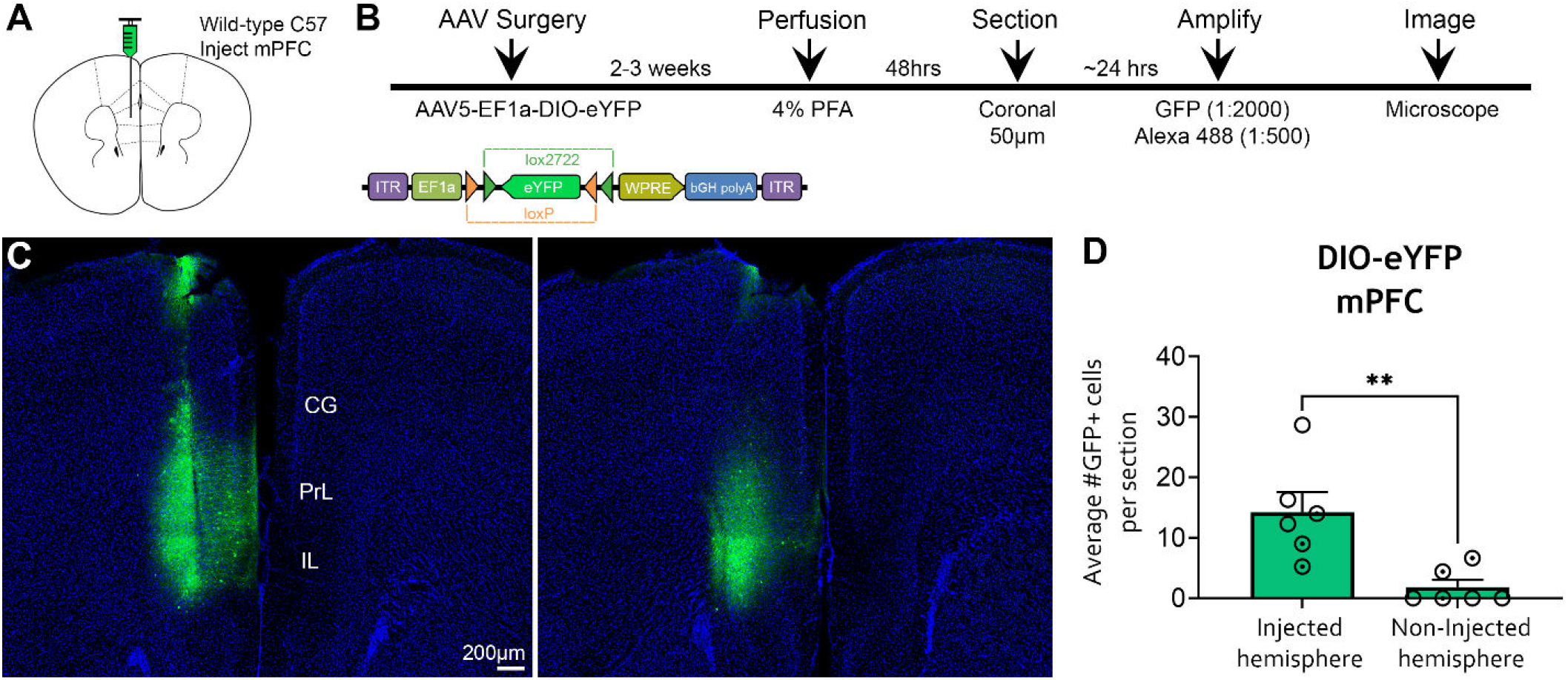
Amplified expression of DIO-eYFP in the mPFC of WT *C57BL/6J* mice. **(A-B)** Experimental design and timeline. AAV5-EF1a-DIO-eYFP was injected into left mPFC of *C57BL/6J* mice (*n*=6) and mice were perfused 2-3 weeks later. Viral signal was amplified with chicken anti-GFP and goat anti-chicken 488 antibodies. **(C)** Representative GFP immunofluorescence in the mPFC of two sections from the same mouse. **(D)** Quantification of GFP+ cells in the mPFC showed that expression was primarily restricted to the injected hemisphere, but two mice had sparse expression of GFP+ cells in the non-injected hemisphere, presumably resulting from viral spread due to the close proximity of the left and right mPFC. Female (clear circles) and male (dotted circles) data points are identified, but no sex differences were found. CG: cingulate gyrus, PrL: prelimbic cortex, IL: infralimbic cortex. **p<0.005. Scale bar: 200µm.

### Fluorescence signal amplification of AAV8-hSyn-DIO-hM3Dq-mCherry in WT *C57BL/6J* mice

To test whether Cre-independent expression with DIO constructs was restricted to a particular AAV serotype, we used AAV8 Cre-dependent hM3Dq (AAV8-hSyn-DIO-hM3Dq-mCherry). Using the same coordinates as eYFP and mCherry experiments described previously, the DIO-hM3Dq construct was injected into the anterior and posterior DG of *C57BL/6* mice (*n*=8; **Figure 6A**). Mice were euthanized 2-3 weeks after surgery and brain sections were processed for mCherry signal amplification (**Figure 6B**). Amplification of AAV8-DIO-hM3Dq-mCherry revealed substantial fluorescence expression in the DG, indicating that Cre-independent expression was observed across multiple serotypes and promoters. Interestingly, amplification of AAV8-DIO-hM3Dq-mCherry construct revealed a different pattern of fluorescence compared with AAV5-DIO-mCherry (**Figure 6C**). Specifically, AAV8-hSyn-DIO-hM3Dq mCherry+ immunofluorescence was primarily observed in hilar neurons, with some sparse labeling in GCs specific to the injected hemisphere (40.73 ± 1.09 cells compared to 0.01 ± 0.01 cells in the non-injected hemisphere; *Mann-Whitney U*=0, *p*<0.001; **Figure 6D**). Notably, the AAV8-hSyn-DIO-hM3Dq-mCherry construct differed from the previous constructs we tested in two ways: serotype (AAV8) and promoter (hSyn, as opposed to EF1a in previous experiments). A subset of sections processed with DAB revealed that mCherry immunoreactivity under the hSyn promoter matched the pattern of amplified DIO-hM3Dq-mCherry immunofluorescence (**Figure 6-1**).

**Figure 6.**
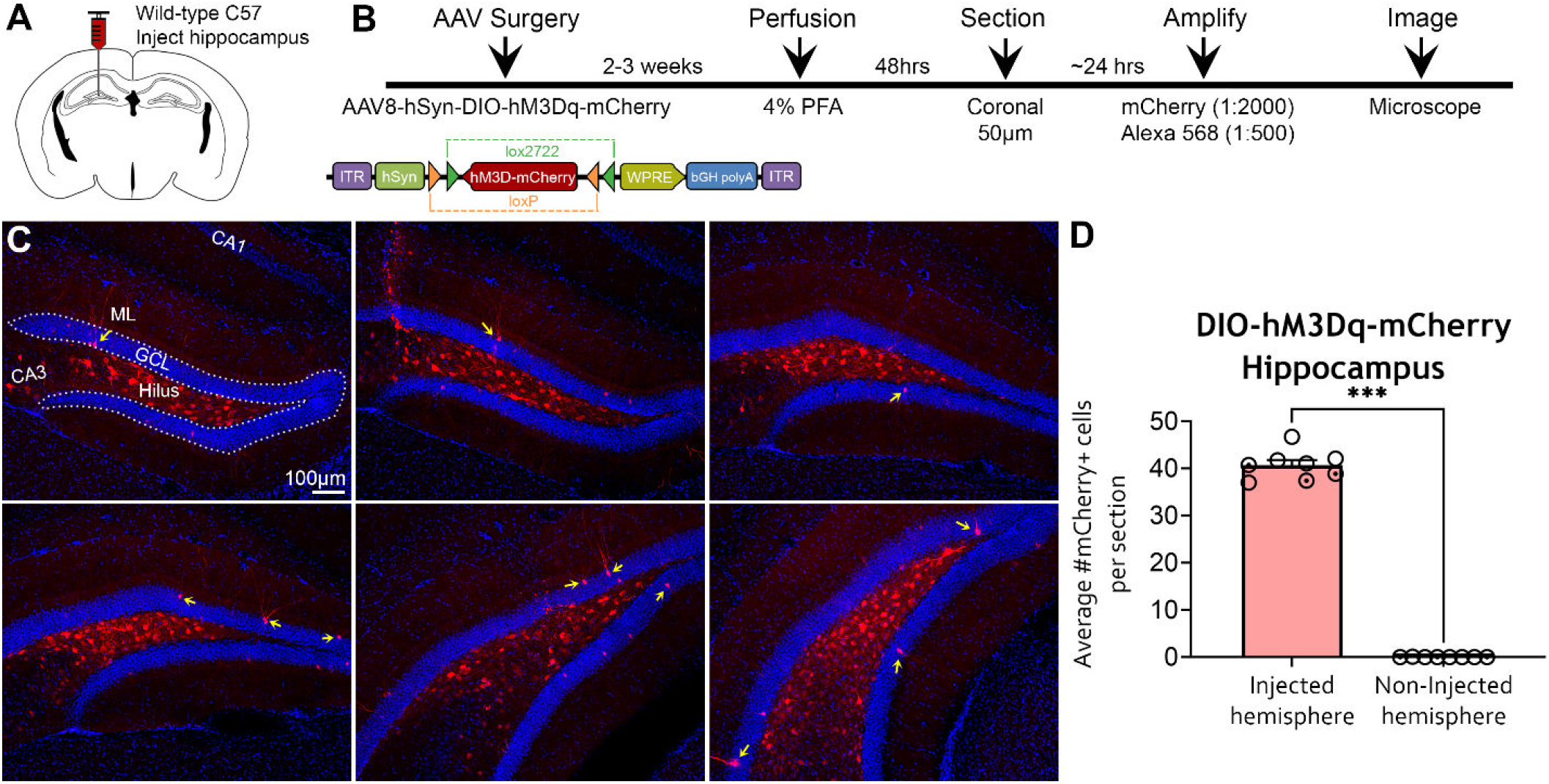
Amplified expression of DIO-hM3Dq-mCherry in the hippocampus of WT *C57BL/6J* mice. **(A-B)** Experimental design and timeline. AAV8-hSyn-DIO-hM3Dq-mCherry was injected into the anterior and posterior hippocampus of *C57BL/6J* mice (*n*=8) and mice were perfused 2-3 weeks later. The viral signal was amplified with rabbit anti-mCherry and goat anti-rabbit 568 antibodies and visualized on an epifluorescence microscope. **(C)** Representative mCherry immunofluorescence in relatively dorsal (top panel) and caudal (bottom panel) sections of the DG. Amplified mCherry expression appeared primarily within hilar cells and a sparse number of GCs (yellow arrows). **(D)** Quantification of mCherry+ cells revealed that expression was restricted to the injected hippocampus. Female (clear circles) and male (dotted circles) data points are identified, but no sex differences were found. GCL: granule cell layer, ML: molecular layer. ***p<0.001. Scale bar: 100µm.

**Figure 6-1.**
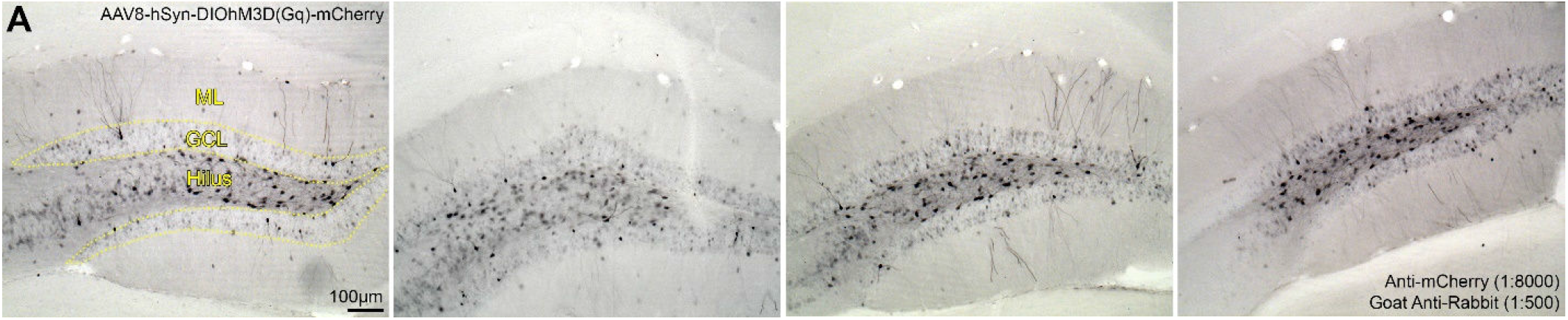
mCherry immunoreactivity in WT *C57BL/6J* mice injected with AAV8-hSyn-DIO-hM3Dq-mCherry. **(A)** Representative photomicrographs of mCherry immunoreactivity in *C57BL/6J* mice injected with AAV8-hSyn-DIO-hM3Dq-mCherry. The pattern of mCherry immunoreactivity was comparable to the amplified immunofluorescence of DIO-hM3Dq-mCherry (see **Figure 6**). GCL: granule cell layer, ML: molecular layer. Scale bar: 100µm.

To determine whether the expression difference was due to serotype, we injected AAV5-hSyn-DIO-hM4Di-mCherry into *C57BL/6J* mice. We found that mCherry amplification of AAV5-hSyn-DIO-hM4Di-mCherry had a similar pattern of fluorescence as AAV8-hSyn-DIO-hM3Dq-mCherry, indicating that serotype is not driving the difference in the pattern of Cre-independent expression (**Figure 6-2**). These results suggest that DIO constructs with the EF1a and hSyn promoters may show preferential expression in GCs vs hilar cells, respectively, in *C57BL/6J* mice. Moreover, these results also demonstrate that off-target expression of DIO constructs was observed using constructs from different vendors (UNC Core, Addgene).

**Figure 6-2.**
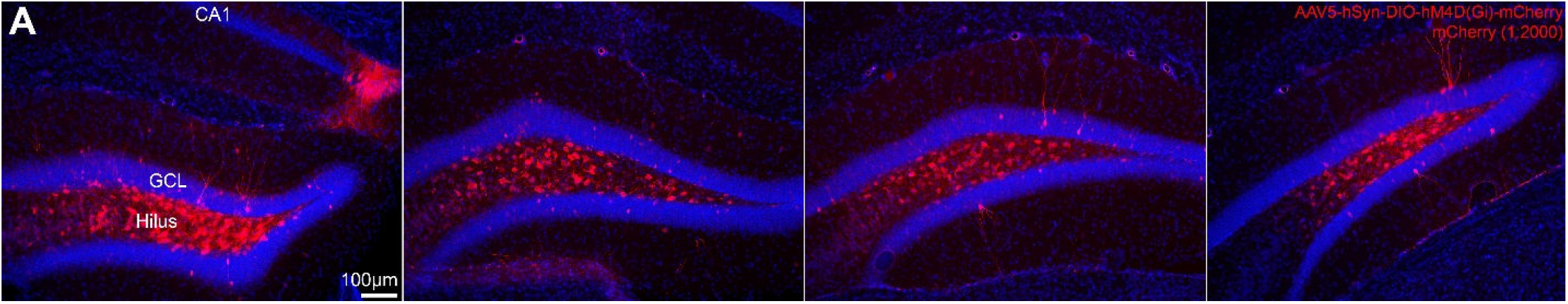
Fluorescence signal amplification of AAV5-hSyn-DIO-hM4Di-mCherry in WT *C57BL/6J* mice. **(A)** *C57BL/6J* mice were injected in the DG with AAV5-hSyn-DIO-hM4D(Gi)-mCherry and sections were amplified with mCherry. Interestingly, mCherry+ cells were primarily located in the hilus, but a small number of GCs were also labeled. The pattern of amplified AAV5-hSyn-DIO-hM4Di-mCherry expression is consistent with the AAV8-hSyn-DIO-hM3Dq-mCherry construct shown in **Figure 6**. Scale bar: 100µm.

### Contextual fear learning and memory

Next, we sought to determine whether the off-target expression of Cre-dependent viral constructs in *C57BL/6J mice* could influence behavior. Given the considerable number of DIO-hM3Dq-mCherry cells observed in the hilus after fluorescence signal amplification (see **Figure 6**), and a recent study that reported chemogenetic excitation of hilar cells impaired contextual fear learning and memory (Botterill et al., 2021), we were curious whether similar impairments would be observed in control mice injected with the DIO construct. Adult *C57BL/6J* mice were injected in the anterior and posterior DG with AAV5-EF1a-DIO-mCherry or AAV8-hSyn-DIO-hM3Dq-mCherry (*n*=8 per group; **Figure 7A**). After a 2-week postsurgical recovery period, mice were injected with the hM3Dq agonist C21 (1.5mg/kg, i.p.) one hour prior to contextual fear training (**Figure 7B-C**).

**Figure 7.**
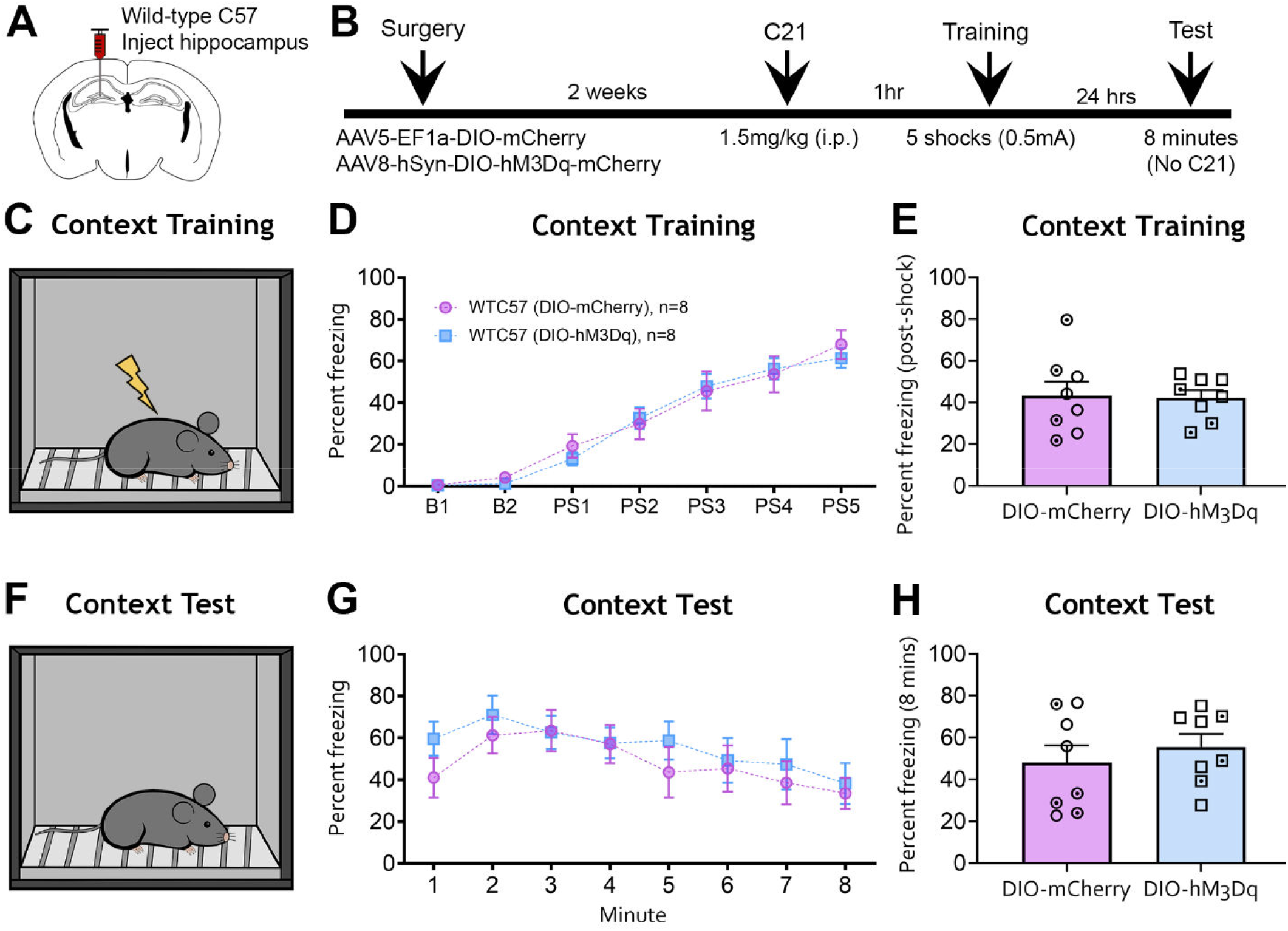
The hM3Dq agonist C21 does not affect fear behavior in *C57BL/6J* mice injected with DIO-mCherry or DIO-hM3Dq-mCherry in the DG. **(A-B)** Experimental design and timeline. Adult *C57BL/6J* mice underwent surgery to receive intrahippocampal injections of AAV-EF1a-DIO-mCherry or AAV-hSyn-DIO-hM3Dq-mCherry. After a 2-week recovery period, mice were injected with the hM3Dq agonist C21 one hour prior to contextual fear training. **(C)** Mice were then placed in a fear conditioning chamber. Baseline activity was assessed over 2 minutes, followed by 5 foot-shocks (0.5mA) spaced 1 minute apart. **(D)** Minute-by-minute analysis of the training session revealed that freezing behavior did not differ between EF1a-DIO-mCherry or hSyn-DIO-hM3Dq-mCherry groups. **(E)** The average post-shock freezing did not differ between the EF1a-DIO-mCherry and hSyn-DIO-hM3Dq-mCherry groups. **(F)** Mice were returned to the same operant chamber 24 hours later to test contextual fear memory. Notably, C21 was not administered a second time prior to the contextual memory test. **(G)** Minute-by-minute analysis revealed that conditioned freezing did not differ between the EF1a-DIO-mCherry or hSyn-DIO-hM3Dq-mCherry groups. **(H)** Average freezing during the memory test did not differ between groups. Female (clear points) and male (dotted points) data points are identified, but no sex differences were found.

C21 treatment prior to contextual fear training had no effect on freezing behavior during training in mice injected with DIO-hM3Dq-mCherry vs DIO-mCherry (Two-way repeated-measures ANOVA, *F*(1,14)=0.045, *p*=0.834; **Figure 7D**). The two-way repeated-measures ANOVA also revealed a significant main effect of time (*F*(6,84)=72.69, *p*<0.001), attributable to increased freezing behavior as the task progressed from baseline freezing to post-shock periods. However, there was no significant interaction between treatment and time (*F*(6,84)=0.474, *p*=0.825). When post-shock freezing was averaged across all 5 post-shock periods, there was no difference in freezing behavior between mice injected with DIO-mCherry (43.32 ± 6.70%) or DIO-hM3Dq-mCherry (42.30 ± 3.63%; unpaired t-test, *t*(14)=0.133, *p*=0.895; **Figure 7E**). Taken together, these results showed no detectable behavioral effect of the hM3Dq agonist C21 in *C57BL/6J* mice injected with DIO-hM3Dq-mCherry.

To evaluate contextual fear memory retrieval, mice were returned to the same fear conditioning chamber 24 hours after training and freezing behavior was evaluated over 8 minutes (**Figure 7F**). Importantly, C21 was not given prior to the memory test. There was no difference in memory retrieval between the DIO-mCherry and DIO-hM3Dq-mCherry groups (Two-way repeated measures ANOVA, *F*(1,14)=0.542, *p*=0.474; **Figure 7G**). However, the two-way repeated-measures ANOVA found a significant main effect of time (*F*(7,98)=4.483, *p*<0.001), which was attributable to a gradual decline in freezing behavior over the duration of the test. There was no interaction between treatment and time (*F*(7,98)=0.512, *p*=0.824). Average freezing behavior over the entire session also did not differ between DIO-mCherry (48.02 ± 8.24%) and DIO-hM3Dq-mCherry groups (55.60 ± 6.15%; unpaired t-test, *t*(14)=0.737, *p*=0.474; **Figure 7H**). Collectively, these results suggest that the hM3Dq agonist C21 did not influence fear learning or memory retrieval in *C57BL/6J* mice injected with DIO-hM3Dq-mCherry.

### mCherry and cFos immunofluorescence following C21 challenge

Despite observing no behavioral effect of C21 in the DIO-hM3Dq-mCherry group, we wanted to determine whether C21 could activate DIO-hM3Dq-mCherry+ neurons in *C57BL/6J* mice by evaluating the immediate early gene cFos. Mice were given a 3-day washout period after fear memory retrieval and then injected with C21 (1.5mg/kg, i.p.) in their homecage and euthanized 90 minutes later (**Figure 8A-B**).

**Figure 8.**
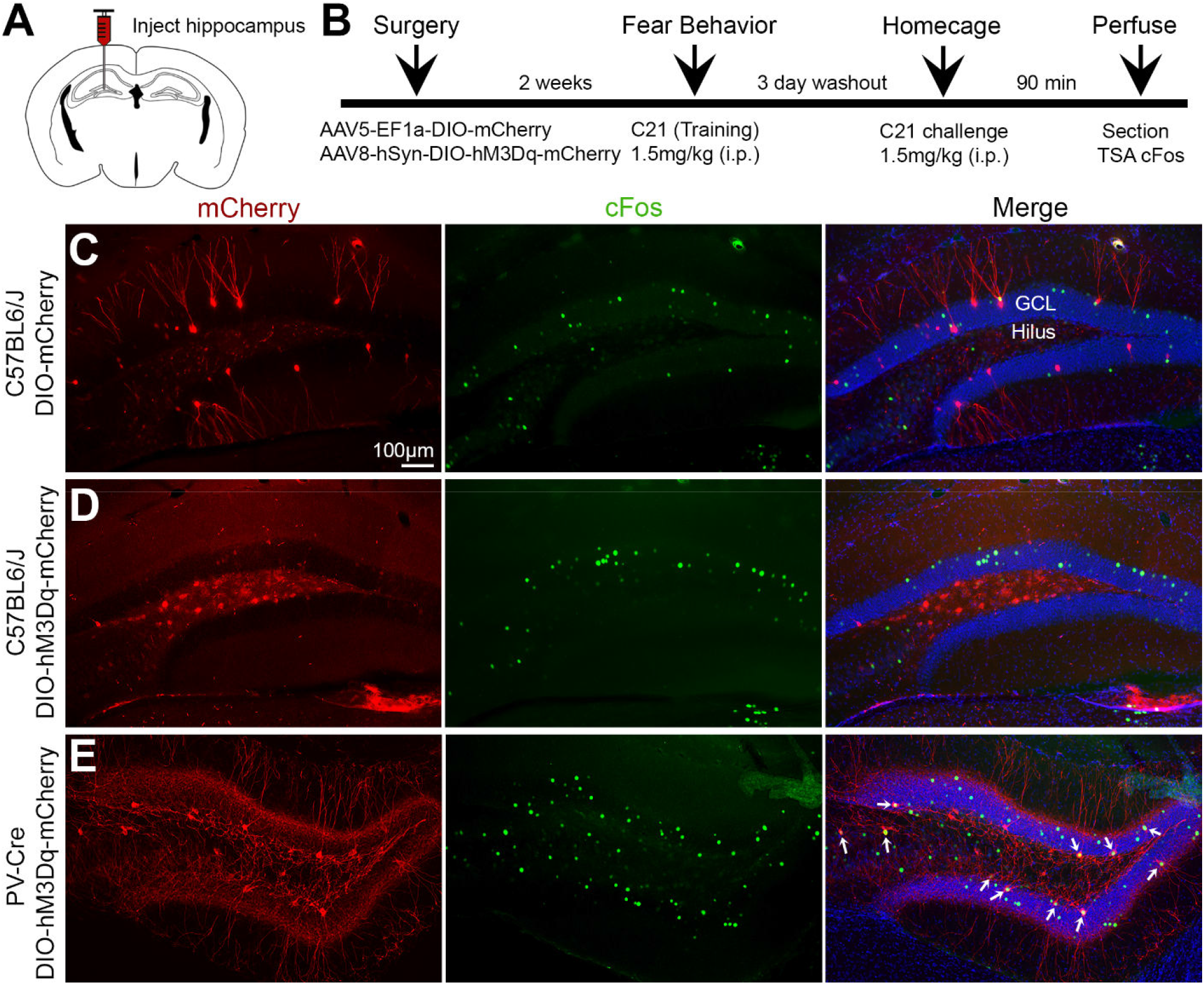
mCherry and cFos immunofluorescence following C21 homecage challenge. **(A-B)** Experimental design and timeline. Mice underwent surgery for AAV injection and allowed 2 weeks to recovery. Mice underwent behavioral testing and were given a 3-day washout period. Mice were then injected with C21 (1.5mg/kg) in their homecage and euthanized 90 minutes later to evaluate the immediate early gene cFos. **(C)** Representative images for C57BL6/J mice injected with DIO-mCherry show no colocalization between amplified mCherry and cFos. **(D)** Representative images for *C57BL/6*J mice injected with DIO-hM3Dq-mCherry show minimal colocalization of cFos and amplified mCherry. **(E)** Representative images for *PV-Cre*-positive mice injected with DIO-hM3Dq-mCherry (red) and challenged with C21 shows a high degree of colocalization with cFos (yellow; white arrows). Scale bar: 100µm.

*C57BL/6J* mice injected with AAV5-EF1a-DIO-mCherry and treated with C21 showed minimal colocalization between amplified mCherry+ cells and cFos+ cells, as expected (**Figure 8C**). Interestingly, when treated with C21, *C57BL/6J* mice injected with AAV8-hSyn-DIO-hM3Dq-mCherry also showed minimal colocalization between mCherry+ and cFos+ cells (**Figure 8D**). This finding suggests that despite DIO-hM3Dq-mCherry expression, the hM3Dq agonist C21 was not able to sufficiently drive their activity as measured by cFos expression. To confirm that DIO-hM3Dq-mCherry is reliably activated by C21 in mice expressing Cre-recombinase, AAV8-hSyn-DIO-hM3Dq-mCherry was injected into a cohort of *PV-Cre*-positive mice (*n*=3) and C21 was administered in the homecage. In contrast to *C57BL/6J* mice, C21 strongly activated DIO-hM3Dq-mCherry+ neurons in the DG of *PV-Cre*-positive mice (**Figure 8E**). Taken together, these results confirm that the hM3Dq agonist C21 potently activates DIO-hM3Dq-mCherry+ neurons in Cre-positive mice, an effect that is absent in WT *C57BL/6J* mice.

## Discussion

The present study investigated anatomical and behavioral effects of Cre-dependent rAAVs in mice lacking Cre-recombinase. WT *C57BL/6J* mice injected with Cre-dependent viral constructs showed minimal non-amplified fluorescence, consistent with the notion that “leak” expression is a rare phenomenon in DIO constructs (Fenno et al., 2011). However, antibody amplification of the fluorescent reporter proteins eYFP or mCherry revealed considerable fluorescence in different brain regions where virus was injected. Subsequent experiments failed to show any behavioral or immediate early gene effect of DIO-hM3Dq-mCherry in *C57BL/6J* mice injected with the hM3Dq agonist C21. These results suggest that Cre-dependent rAAVs injected in mice lacking Cre can result in off-target transgene expression, as revealed by fluorescence signal after antibody amplification, but without yielding notable behavioral or functional effects in our experimental system.

### Fluorescence signal amplification of viral expression

In this work we evaluated fluorescence signal amplification in Cre-positive and *C57BL/6J* mice injected with various Cre-dependent rAAVs. First, we evaluated *TH-Cre*-positive mice injected with DIO-mCherry and found that the expression of fluorescently labeled cell bodies in the VTA were consistent with previous studies (Stuber et al., 2010; Mahler et al., 2019). However, non-amplified fluorescence of VTA projections into the NAc-DS or mPFC were notably weak and fluorescence signal amplification improved the visualization of mCherry, especially in mPFC axon terminals (see **Figure 1C**). We also evaluated *PV-Cre*-positive mice injected with DIO-mCherry or DIO-eYFP in the DG and found that while non-amplified fluorescence was suitable for visualizing PV+ cells, fluorescence signal amplification improved expression in fine processes such as dendrites. Collectively, these findings support the notion that fluorescence signal amplification can significantly improve visualization of viral transgene expression (Falcy et al., 2020).

We also tested the specificity of Cre-dependent rAAVs in *C57BL/6J* mice. We observed minimal non-amplified fluorescence, consistent with the dependence of Cre-recombinase to drive transgene expression (Fenno et al., 2011; Fischer et al., 2019). However, we found that fluorescence signal amplification reliably labeled mCherry+ or GFP+ cells wherever the Cre-dependent rAAV was injected (e.g., DG, CA1 or mPFC). Importantly, there were few or no amplified cells in the non-injected hemisphere, suggesting that antibody specificity was not an issue. Furthermore, fluorescence amplification revealed substantial AAV-DIO expression in *C57BL/6J* mice regardless of the commercial vendor (Addgene, UNC Core), serotype (AAV5, AAV8) or promoter (EF1a, hSyn) used. These observations indicate that our results could apply to a broad range of rAAV DIO constructs. Overall, these findings warrant caution in interpreting the results of DIO constructs in Cre-negative subjects, especially if quantitative measures are used following fluorescence signal amplification.

### Functional considerations

Upon discovering the effect of fluorescence signal amplification in *C57BL/6J* mice injected with Cre-dependent rAAVs, we considered the implications for off-target effects in subjects typically assigned as controls. We used the hM3Dq agonist C21 to determine whether the expression of DIO-hM3Dq in *C57BL/6J* mice had any functional effects. We found that C21 had no impact on contextual fear learning or memory retrieval and was insufficient to trigger cFos expression in DIO-hM3Dq-mCherry+ cells in *C57BL/6J* mice. In contrast, C21 induced cFos expression in DIO-hM3Dq-mCherry+ cells of *PV-Cre*-positive mice. Our results are consistent with previous studies that found no effect of DIO-hM3Dq in Cre-negative subjects injected with DREADD agonists compared to Cre-positive counterparts (Alexander et al., 2018; Bonaventura et al., 2019; Mahler et al., 2019), suggesting that DIO-construct expression levels in *C57BL/6J* mice may be insufficient to modulate neuronal activity and affect behavior. Nevertheless, expression level thresholds for phenotypic change will differ between experimental contexts, and as such it cannot be ruled out that functional consequences can arise from off-target gene expression from Cre-dependent rAAV.

### Technical Considerations Viral titer and injection volume

Specificity of viral expression is a common concern in experiments that use rAAVs. Viral titer and injection volume represent two main factors that can impact viral expression, and thereby might modulate DIO-construct expression in Cre-negative animals. High titer rAAVs are required to introduce numerous viral particles within a single cell to achieve adequate viral expression.

For neuroscience applications, commercial vendors typically provide rAAVs at titers ranging between ≥1 × 10^11^vg/mL to ∼10^13^vg/mL. However, the relationship between vector dose and protein expression is non-linear. For example, a study reported a 6-fold increase in the number of virally labeled cells when viral titer was adjusted from 5×10^12^vg/mL to 5×10^13^vg/mL (Zingg et al., 2017). A second factor to consider is viral injection volume, which is often influenced by factors such as experimental design or the size of the brain region that is targeted. For large brain regions like the hippocampus, injection volumes of ∼0.25µL are relatively common, but numerous studies have injected volumes ≥0.5µL and report good specificity (Gundersen et al., 2013; Bui et al., 2018; Piatkevich et al., 2019; Johnston et al., 2021).

In the present study, stock rAAV titers from commercial vendors (≥4 × 10^12^vg/mL) were used at relatively low injection volumes (0.2µL) because these parameters achieved highly specific expression in *TH-Cre*-positive and *PV-Cre*-positive mice. In *C57BL/6J* mice, this injection volume yielded minimal non-amplified fluorescence, but considerable immunofluorescence following antibody amplification. Dilution of viral titer has been proposed as a mitigation strategy to minimize off-target rAAV expression; however, dose reduction could potentially have a negative impact on experimental outcomes by missing phenotypes that are only observable with robust transgene expression.

### Causes of off-target expression in mice lacking Cre

The cause of off-target Cre-independent rAAV transgene expression was not investigated within the scope of this study. Spontaneous reversion of DIO constructs is known to occur at a low rate and is likely to be the origin of some of this expression. In support of this, a previous study evaluated recombinant plasmids and found that between 1 in 1,000 and 1 in 10,000 copies contained a reverted transgene (Fischer et al., 2019).

However, given our detection of substantial numbers of low intensity transgene expressing cells, we suspect that there are factors additional to transgene reversion that could result in Cre-independent expression of DIO constructs. The ITRs of AAV are known to exhibit transcriptional activity in a number of cell types, with the AAV2 ITRs, used in the majority of applications, exhibiting stronger promoter activity than ITRs from several other serotypes (Earley et al., 2020). Indeed, early rAAV gene therapy constructs for cystic fibrosis relied on this activity to drive expression of the CFTR gene (Flotte et al., 1992). It is possible that in *C57BL/6J* mice, weak expression of the transgene could be achieved through transcriptional activity of the ITR, although transcriptional activity of ITRs is yet to be directly tested in neuronal cell populations.

Furthermore, within the nucleus, rAAV largely exists in a concatemeric, episomal state (Yang et al., 1999). Where the head is the 5’ end of the rAAV genome and the tail is the 3’, the configurations of multiple rAAV genomes can either be head-to-head, head-to-tail or tail-to-tail. If multiple copies of non-reverted DIO constructs were present within a single cell, it is possible that in the tail-to-tail configuration, promoter activity from one DIO genome could readthrough the rAAV sequence to translate the encoded protein in a second genome of the concatemer. Indeed, this reliance on transcription across multiple genomes is used to yield expression from large gene constructs, using splice donor and acceptor sites in the two respective rAAV genomes (Trapani et al., 2015).

Finally, whilst AAV is largely considered to be a non-integrating vector, it is known that integration events do occur at low levels. It is possible that if the DIO construct integrated at a transcriptionally active locus, translation of the non-reverted transgene could be initiated. Indeed, this is the basis which promoterless rAAV constructs for rAAV-mediated gene therapy operate, albeit in a more actively targeted and efficient manner (Barzel et al., 2015).

### Minimizing off-target expression in DIO constructs

A previous study revealed that both loxP site mutation and decoupling the start codon from the gene to a position outside of the loxP inversion sites were required to achieve dramatic reduction in off-target expression from DIO/FLEX rAAV constructs, a system referred to as ‘ATG-out’ (Fischer et al., 2019). This suggests that transgene reversion is not the only cause of off-target expression in neurons following DIO construct delivery, because if this was the case, loxP mutation alone would have been sufficient to minimize this effect. At present, this strategy has not been widely implemented in the neuroscience field, but should be considered by those using sensitive systems and/or cell counting assays. Importantly, the ATG-out system, whilst vastly reducing off-target activity, did not entirely abrogate expression in the system, and was not assessed within the context of signal amplification. Further work should be performed to ensure the fidelity of ATG-out vectors in signal amplified samples, and to explore other approaches for improving the specificity of inducible transgene systems for use in neuroscience applications.

### Specificity of Cre-recombinase

Cre-dependent rAAVs are generally considered to have a high degree of specificity due to the dependance of Cre-recombinase to drive transgene expression (Huang et al., 2014; Saunders and Sabatini, 2015; McLellan et al., 2017; Haggerty et al., 2020). However, specificity of Cre-recombinase can be influenced by factors such as breeding, genotyping, and/or germline recombination (Song and Palmiter, 2018). Specificity problems are particularly well-documented in tamoxifen-inducible transgenic lines (Stifter and Greter, 2020; Van Hove et al., 2020). Therefore, it is important to consider the specificity of transgenic lines in addition to rAAV titer and injection volume.

### Implications for control experiments

Selecting appropriate controls is a critical step in designing rAAV experiments, especially for studies that involve cell and/or circuit manipulations. There are several strategies for rAAV controls, and each approach has strengths and weaknesses. For example, a popular strategy involves injecting Cre-positive mice with identical rAAV constructs and randomly assigning subjects to a treatment (e.g., CNO or C21) or control group (e.g., saline). Although this strategy controls for genotype and viral construct, it often overlooks the effect of treatment. Indeed, compounds such as CNO can have off-target effects (MacLaren et al., 2016; Gomez et al., 2017; Manvich et al., 2018) and therefore these experiments often require additional controls that receive treatment but not the same rAAV construct. A second strategy involves comparing Cre-positive vs Cre-negative littermates injected with identical rAAV constructs (Smith et al., 2016). This strategy offers the benefit of treating all subjects identically but does not account for potential genotype effects in Cre-positive mice. Moreover, this strategy requires additional steps such as confirmation of genotypes and/or evaluation of viral expression in Cre-positive vs Cre-negative mice. Lastly, another popular strategy involves injecting Cre-positive mice with gain- or loss-of function rAAV constructs and control mice with an rAAV construct that only encodes a fluorescent protein such as mCherry or eYFP. This strategy also allows for all mice to receive the same treatment (e.g., CNO or light pulses). This approach is widely used because of the low risk of off-target effects in control mice, but the disadvantage is the use of different viral constructs.

Although we did not observe any functional off-target effects of Cre-dependent rAAVs in the DG of *C57BL/6J* mice, we did not evaluate factors such as different behavioral tasks, greater rAAV injection volumes (e.g., 0.5µL), rAAV injections in different brain regions, or higher doses of C21. Based on the results of the current study, we suggest caution when choosing controls for gain- or loss-of function Cre-dependent constructs. Our data points to the use of fluorophore-only controls as the preferential option to minimize potential off-target effects of Cre-dependent rAAV constructs in control mice.

## Conclusions

Cre-recombinase dependent rAAVs represent a powerful tool that many neuroscientists utilize for labeling, tracing, or manipulating specific neuronal populations. Although the fluorescent reporter of most viral constructs yields suitable transgene expression levels within infected cell populations, many laboratories utilize antibody-based fluorescence signal amplification to visualize weak or intermediate fluorescence signals. Here, we report the observation that Cre-dependent AAVs injected into different brain regions of mice lacking Cre-recombinase reliably showed expression following antibody amplification of the fluorescent reporter. Our results therefore caution that researchers must carefully design and interpret data involving Cre-dependent rAAV infection.

## Acknowledgements

We thank Drs. David Alcantara-Gonzalez and Vinod Yaragudri for their contributions at the early stages of this project. We thank Dr. Jonathan Britt for sending us the *TH-Cre* mice.

